# Bidirectional Mendelian randomization analysis of shared genetic signals between coexisting neurodegenerative disorders to decipher underlying causal pathways

**DOI:** 10.1101/692921

**Authors:** Sandeep Grover, International Age-related Macular Degeneration Consortium (IAMDGC)

## Abstract

**OBJECTIVE:** To investigate whether coexistence of various neurodegenerative disorders is coincidental or biologically connected.

**DESIGN:** Two sample Mendelian randomization using summary effect estimates

**SETTING:** Genetic data taken on various neurodegenerative disorders from various cohorts comprising individuals predominantly of European ancestry.

**PARTICIPANTS:** International Genomics of Alzheimer’s patients (IGAP), project MinE, International Age-related Macular Degeneration Consortium (IAMDGC), International Multiple Sclerosis Genetics Consortium (IMSGC), International Parkinson’s Disease Genomics Consortium (IPDGC)

**MAIN OUTCOME MEASURES:** Alzheimer’s disease (AD), Amyotrophic lateral sclerosis (ALS), Age related macular degeneration (AMD), Multiple sclerosis (MS) and Parkinson’s disease (PD).

**RESULTS:** A Bonferroni corrected threshold of P=0.005 was considered to be significant, and P<0.05 was considered suggestive of evidence for a potential association. I observed a risky effect of PD on ALS (OR = 1.126, 95% CI = 1.059-1.198, P = 0.005). Using AD as exposure and PD as outcome, I observed a risky effect of AD on PD using all the MR methods with strongest results using MBE method (OR = 2.072, 95% CI = 1.006-4.028, P = 0.0416). Genetic predisposition to AD was further observed to be a risky for AMD (OR = 1.759, 95% CI = 1.040-1.974, P = 0.0363). On the contrary, AMD was observed to be strongly protective towards MS (OR = 0.861, 95% CI = 0.776-0.955, P = 0.0059).

**CONCLUSIONS:** My findings are consistent with the previously observed relative occurrence of co-existing neurodegenerative diseases or overlapping symptoms among neurodegenerative diseases.

## Introduction

It is not uncommon to see cases of neurodegeneration in clinical practice showing temporal development of one neurodegenerative disorder after another. For instance, observational studies have shown higher risk of Parkinson’s disease (PD) and Alzheimers’ disease (AD) among patients with a diagnosis of neovascular age-related macular degeneration (AMD)^1–4^. Occasional case reports of co-existence of Multiple sclerosis (MS) with Alzheimer’s disease (AD) have also emerged in the literature^5^. Furthermore, the co-existence of Amyotrophic lateral sclerosis (ALS) with Frontotemporal dementia (FTD) had recently lead to revises consensus criteria for the diagnosis of FTD in ALS^6^.

Not surprisingly, most neurodegenerative diseases share certain clinical and pathological features. Furthermore, a number of genetic studies have time and again have also shown existence of shared genetic aetiology^7–9^. It is common to see overlapping symptoms among various neurodegenerative disorders. For instance, up to 50% of AD cases exhibit aggregation of alpha-synuclein into Lewy bodies, a characteristic seen in PD cases^10^. Furthermore, degeneration of retinal layer, a characteristic of AMD has also been reported in cases with ALS and MS^11, 12^. Of all the co-existing neurodegenerative, presence of ALS, parkinsonism and dementia together is the most well characterized combination often reported in specific geographic locations and known by several name such as kii-ALS or Guam-ALS or ALS-PDC^13, 14^.

It has been long debated whether co-existence of neurodegenerative disorders is purely coincidental or there is a causal relationship in between them. However, the varying latent phases of different neurodegenerative disorders make it difficult to interpret the exact relationship between the co-existing disorders. With age as a major confounding factor in observational studies, Mendelian randomization (MR) methodology could provide an alternative solution by providing life-long effect estimates using genetic variants as proxy pseudorandomized markers of neurodegenerative diseases^15^. Henceforth, the objective of the current study was to explore the causal relationships among different neurodegenerative disorders using MR approach.

## Methods

A two-sample MR approach was applied to explore the relationship among six most commonly occurring neurodegenerative disorders namely Alzheimer’s disease (AD), Amyotrophic lateral sclerosis (ALS), Age-related macular degeneration (AMD), Frontotemporal dementia (FTD), Multiple sclerosis (MS) and Parkinson’s disease (PD)^16–20^. I employed latest available summary GWAS datasets for the present study to prioritize genetic instruments for each of the neurodegenerative disorder. I used inverse variance weighted method as the main method to generate unconfounded estimates using the summary statistics from respective GWAS datasets to explore the relationship between each pair of neurodegenerative disorder. A Bonferroni corrected threshold of P=0.005 (as ten pairs of neurodegernative disorders were compared) was considered to be significant, and P<0.05 was considered suggestive of evidence for a potential association.

I also generated causal estimates adjusted for presence of potential pleiotropic variants by employing additional MR methods. The heterogeneity in the effect estimates were judged using MR-Egger, I2 and Cochrane-Q statistics. Lastly, sensitivity analysis was conducted to check reliability of estimates by excluding variants known to be directly involved in specific neurodegenerative disorder as an outcome, and variants believed to be potential confounders between pair of neurodegenerative disorder under consideration.

## Results

All the datasets used in the study have been shown in **Table 1**. In addition, complete summary statistics used for causal analysis is provided in **Supplementary table 1**. The results from direct and reverse causal estimate analysis have been provided in **Table 2a to 2e**. I observed a risky effect of PD on ALS (OR = 1.126, 95% CI = 1.059-1.198, P = 0.005). The risky effect was further retained using IVW, MR-Egger and WME method. Furthermore, a weak bidirectional relationship was observed between AD and PD. Using AD as exposure and PD as outcome, I observed a risky effect of AD on PD using all the MR methods with strongest results using MBE method (OR = 2.072, 95% CI = 1.006-4.028, P = 0.0416). A moderate risky effect of PD on AD was further observed using WME method (OR = 1.013, 95% CI = 1.006-1.019, P = 0.0606).

**Table 1.** Details of discovery GWAS datasets explored and prioritized instruments used for the main analysis in the present study.

| S.No. | Exposure | Source study | Maximum sample size | GWAS study cohort | Number of SNPs analyzed | P | Number of significant SNPs (pre-clumping) | Number of significant SNPs (post-clumping) | Average F-statistics (median (range)) |
| --- | --- | --- | --- | --- | --- | --- | --- | --- | --- |
| 1 | Alzheimer's disease (AD) (AD) | Jansen et al. 2018 | 71880 cases/ 383,378 controls | Discovery | 133,672,99 | $5 \times 10^{-8}$ | 2357 | 26 | 42.2 (30.2-422.5) |
| 2 | Amyotrophic lateral sclerosis (ALS) | van Rheenen et al. 2016 | 12577 cases/ 23475 controls | Discovery | 870, 945,2 | $5 \times 10^{-8}$ | 125 | 4 | 37.2 (32.2-80.1) |
| 3 | Age-related macular degeneration (AMD) | Fritsche et al. 2016 | 16144 cases/17832 controls | Discovery | 120,238,39 | $5 \times 10^{-8}$ | 7218 | 42 | 47.5 (29.2-382.5) |
| 4 | Multiple sclerosis (MS) (MS) | Patsopoulos et al. 2017 | 47351 cases/ 68284 controls | Discovery | 859, 365,0 | $5 \times 10^{-8}$ | 26403 | 74 | 41.9 (29.8-561.9) |
| 5 | Parkinson's disease (PD) (PD) | Nalls et al. 2018 | 33,674 cases/ 449, 056 controls | Discovery | 175,137,73 | $5 \times 10^{-8}$ | 3465 | 23 | 43.6 (30.0-181.5) |

\*Clumping was done using TwoSampleMR R package

**Table 2a.**
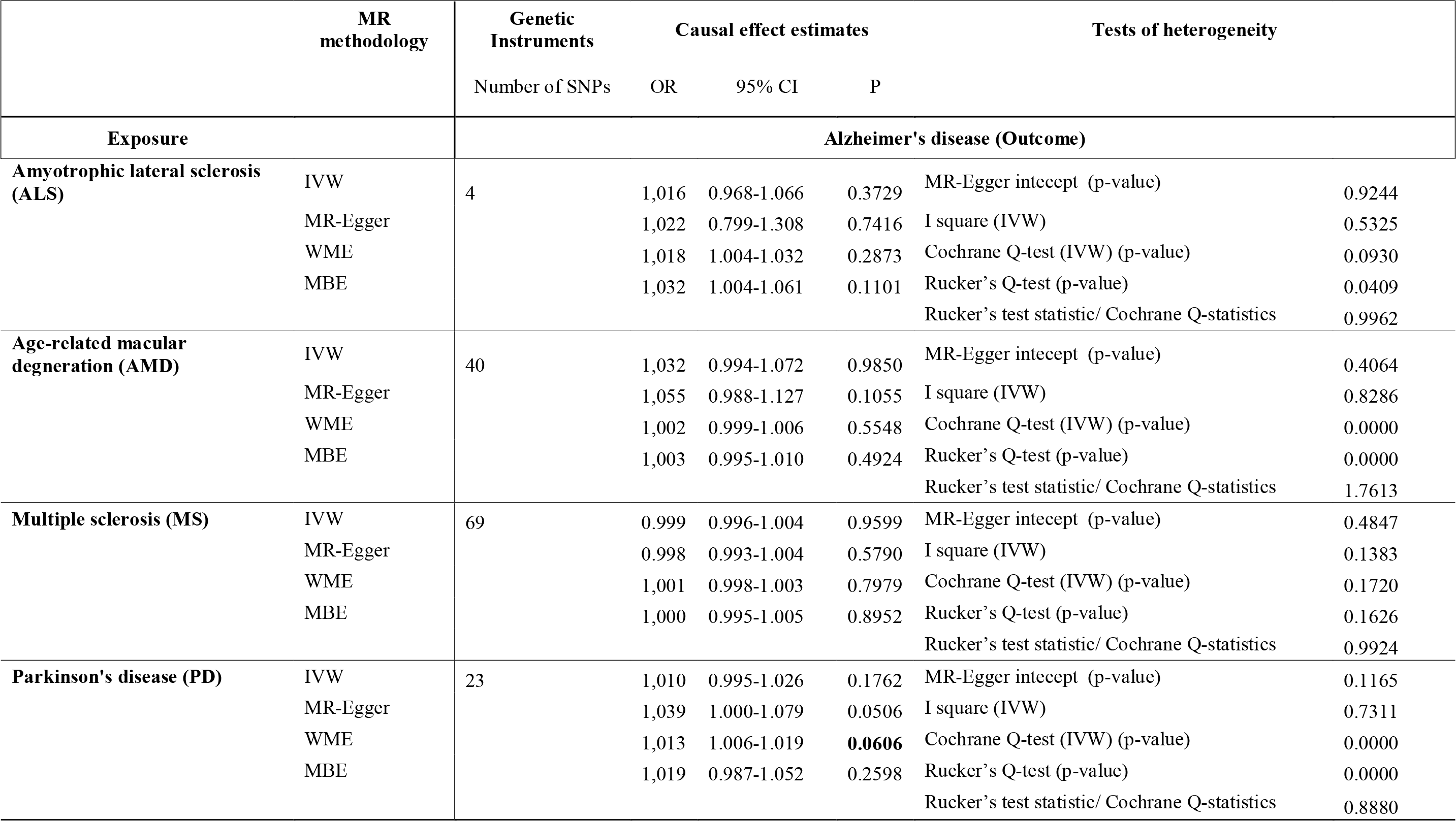
Causal effect estimates exploring influence of various neurodegenerative disorders on Alzheimer’s disaease (AD).

**Table 2b.** Causal effect estimates exploring influence of various neurodegenerative disorders on Amyotrophic lateral sclerosis (ALS).

|  | MR methodology | Genetic Instruments | Causal effect estimates |  |  | Tests of heterogeneity |  |
| --- | --- | --- | --- | --- | --- | --- | --- |
|  |  | Number of SNPs | OR | 95% CI | P |  |  |
| Exposure |  | Amyotrophic lateral sclerosis (ALS) (Outcome) |  |  |  |  |  |
| Alzheimer's disease (AD) | IVW | 25 | 0.996 | 0.623-1.591 | 0.9849 | MR-Egger intercept (p-value) | 0.3992 |
|  | MR-Egger |  | 1,729 | 0.422-7.083 | 0.4299 | I square (IVW) | 0.0000 |
|  | WME |  | 1,327 | 0.948-1.857 | 0.4087 | Cochrane Q-test (IVW) (p-value) | 0.6550 |
|  | MBE |  | 1,584 | 0.497-5.043 | 0.4443 | Rucker's Q-test (p-value) | 0.6395 |
|  |  |  |  |  |  | Rucker's test statistic/ Cochrane Q-statistics | 0.9670 |
| Age-related macular degeneration (AMD) | IVW | 42 | 0.986 | 0.945-1.030 | 0.5229 | MR-Egger intercept (p-value) | 0.4077 |
|  | MR-Egger |  | 1,012 | 0.938-1.092 | 0.7495 | I square (IVW) | 0.1823 |
|  | WME |  | 1,002 | 0.973-1.032 | 0.9539 | Cochrane Q-test (IVW) (p-value) | 0.1550 |
|  | MBE |  | 1,015 | 0.948-1.086 | 0.6724 | Rucker's Q-test (p-value) | 0.1459 |
|  |  |  |  |  |  | Rucker's test statistic/ Cochrane Q-statistics | 0.9857 |
| Multiple sclerosis (MS) | IVW | 72 | 1,008 | 0.984-1.033 | 0.4934 | MR-Egger intercept (p-value) | 0.3323 |
|  | MR-Egger |  | 1,020 | 0.986-1.054 | 0.2464 | I square (IVW) | 0.0269 |
|  | WME |  | 1,034 | 1.015-1.053 | 0.0786 | Cochrane Q-test (IVW) (p-value) | 0.4134 |
|  | MBE |  | 1,023 | 0.992-1.056 | 0.1566 | Rucker's Q-test (p-value) | 0.4151 |
|  |  |  |  |  |  | Rucker's test statistic/ Cochrane Q-statistics |  |
| Parkinson's disease (PD) | IVW | 22 | 1,126 | 1.059-1.198 | **0.0006** | MR-Egger intercept (p-value) | 0.1318 |
|  | MR-Egger |  | 1,280 | 1.068-1.533 | **0.0099** | I square (IVW) | 0.0000 |
|  | WME |  | 1,111 | 1.064-1.161 | **0.0239** | Cochrane Q-test (IVW) (p-value) | 0.6965 |
|  | MBE |  | 1,083 | 0.928-1.264 | 0.3248 | Rucker's Q-test (p-value) | 0.7844 |
|  |  |  |  |  |  | Rucker's test statistic/ Cochrane Q-statistics | 0.8619 |

**Table 2c.** Causal effect estimates exploring influence of various neurodegenerative disorders on Age-related macular degeneration (AMD).

|  | MR methodology | Genetic Instruments | Causal effect estimates |  |  | Tests of heterogeneity |  |
| --- | --- | --- | --- | --- | --- | --- | --- |
|  |  | Number of SNPs | OR | 95% CI | P |  |  |
| Exposure |  | Age-related macular degeneration (AMD) (Outcome) |  |  |  |  |  |
| Alzheimer's disease (AD) | IVW | 26 | 1,759 | 1.040-2.974 | **0.0363** | MR-Egger intecept (p-value) | 0.2093 |
|  | MR-Egger |  | 3,150 | 1.083-9.162 | **0.0362** | I square (IVW) | 0.3513 |
|  | WME |  | 1,678 | 1.238-2.275 | 0.1010 | Cochrane Q-test (IVW) (p-value) | 0.0410 |
|  | MBE |  | 1,683 | 0.791-3.581 | 0.1886 | Rucker's Q-test (p-value) | 0.0571 |
|  |  |  |  |  |  | Rucker's test statistic/ Cochrane Q-statistics | 0.9295 |
| Amyotrophic lateral sclerosis (ALS) | IVW | 4 | 0.924 | 0.679-1.256 | 0.4717 | MR-Egger intecept (p-value) | 0.4079 |
|  | MR-Egger |  | 1,209 | 0.369-3.959 | 0.5629 | I square (IVW) | 0.3516 |
|  | WME |  | 0.985 | 0.900-1.078 | 0.8782 | Cochrane Q-test (IVW) (p-value) | 0.2013 |
|  | MBE |  | 1,015 | 0.841-1.225 | 0.8879 | Rucker's Q-test (p-value) | 0.2119 |
|  |  |  |  |  |  | Rucker's test statistic/ Cochrane Q-statistics | 0.6708 |
| Multiple sclerosis (MS) | IVW | 68 | 0.976 | 0.939-1.015 | 0.2239 | MR-Egger intecept (p-value) | 0.9725 |
|  | MR-Egger |  | 0.978 | 0.870-1.099 | 0.7054 | I square (IVW) | 0.2083 |
|  | WME |  | 0.968 | 0.943-0.993 | 0.2107 | Cochrane Q-test (IVW) (p-value) | 0.0717 |
|  | MBE |  | 0.932 | 0.817-1.064 | 0.3023 | Rucker's Q-test (p-value) | 0.0608 |
|  |  |  |  |  |  | Rucker's test statistic/ Cochrane Q-statistics | 1.0002 |
| Parkinson's disease (PD) | IVW | 23 | 0.953 | 0.901-1.007 | 0.0863 | MR-Egger intecept (p-value) | 0.7136 |
|  | MR-Egger |  | 0.931 | 0.806-1.074 | 0.3098 | I square (IVW) | 0.0000 |
|  | WME |  | 0.929 | 0.896-0.963 | **0.0556** | Cochrane Q-test (IVW) (p-value) | 0.7723 |
|  | MBE |  | 0.927 | 0.836-1.028 | 0.1668 | Rucker's Q-test (p-value) | 0.7281 |
|  |  |  |  |  |  | Cochrane Q-staitics/Rucker's test statistic | 0.9929 |

**Table 2d.** Causal effect estimates exploring influence of various neurodegenerative disorders on Multiple sclerosis (MS).

|  | MR methodology | Genetic Instruments | Causal effect estimates |  |  | Tests of heterogeneity |  |
| --- | --- | --- | --- | --- | --- | --- | --- |
|  |  | Number of SNPs | OR | 95% CI | P |  |  |
| **Exposure** |  | **Multiple sclerosis (MS) (Outcome)** |  |  |  |  |  |
| **Alzheimer's disease (AD)** | IVW | 26 | 1,597 | 0.257-9.908 | 0.6020 | MR-Egger intecept (p-value) | 0.7683 |
|  | MR-Egger |  | 3,032 | 0.025-374.146 | 0.6388 | I square (IVW) | 0.7059 |
|  | WME |  | 1,305 | 0.917-1.857 | 0.4577 | Cochrane Q-test (IVW) (p-value) | 0.0000 |
|  | MBE |  | 1,445 | 0.563-3.708 | 0.4508 | Rucker's Q-test (p-value) | 0.0000 |
|  |  |  |  |  |  | Rucker's test statistic/ Cochrane Q-statistics | 1.0094 |
| **Amyotrophic lateral sclerosis (ALS)** | IVW | 3 | 0.998 | 0.629-1.582 | 0.9860 | MR-Egger intecept (p-value) | 0.3454 |
|  | MR-Egger |  | 1,822 | 0.016-204.153 | 0.3528 | I square (IVW) | 0.4493 |
|  | WME |  | 0.976 | 0.888-1.072 | 0.8186 | Cochrane Q-test (IVW) (p-value) | 0.1627 |
|  | MBE |  | 1,142 | 0.940-1.388 | 0.3138 | Rucker's Q-test (p-value) | 0.3285 |
|  |  |  |  |  |  | Rucker's test statistic/ Cochrane Q-statistics | 0.2629 |
| **Age-related macular degeneration (AMD)** | IVW | 35 | **0.939** | **0.884-0.997** | **0.0409** | MR-Egger intecept (p-value) | 0.0481 |
|  | MR-Egger |  | **0.861** | **0.776-0.955** | **0.0059** | I square (IVW) | 0.4715 |
|  | WME |  | 0.966 | 0.930-1.004 | 0.3816 | Cochrane Q-test (IVW) (p-value) | 0.0013 |
|  | MBE |  | 1,065 | 0.955-1.187 | 0.2652 | Rucker's Q-test (p-value) | 0.0055 |
|  |  |  |  |  |  | Cochrane Q-statistics/Rucker's test statistic | 0.8898 |
| **Parkinson's disease (PD)** | IVW | 22 | 1,042 | 0.976-1.112 | 0.2030 | MR-Egger intecept (p-value) | 0.5749 |
|  | MR-Egger |  | 1,087 | 0.918-1.287 | 0.3145 | I square (IVW) | 0.1214 |
|  | WME |  | 1,049 | 1.005-1.095 | 0.2722 | Cochrane Q-test (IVW) (p-value) | 0.2978 |
|  | MBE |  | 1,004 | 0.863-1.167 | 0.9627 | Rucker's Q-test (p-value) | 0.2655 |
|  |  |  |  |  |  | Rucker's test statistic/ Cochrane Q-statistics | 0.9826 |

**Table 2e.** Causal effect estimates exploring influence of various neurodegenerative disorders on Parkinson’s disease (PD).

|  | MR methodology | Genetic Instruments | Causal effect estimates |  |  | Tests of heterogeneity |  |
| --- | --- | --- | --- | --- | --- | --- | --- |
|  |  | Number of SNPs | OR | 95% CI | P |  |  |
| **Exposure** |  | **Parkinson's disease (PD) (Outcome)** |  |  |  |  |  |
| **Alzheimer's disease (AD)** | IVW | 26 | 2,189 | 0.996-4.812 | **0.0510** | MR-Egger intecept (p-value) | 0.3779 |
|  | MR-Egger |  | 1,270 | 0.289-5.584 | 0.7418 | I square (IVW) | 0.5830 |
|  | WME |  | 1,951 | 1.415-2.688 | **0.0477** | Cochrane Q-test (IVW) (p-value) | 0.0001 |
|  | MBE |  | 2,072 | 1.066-4.028 | **0.0416** | Rucker's Q-test (p-value) | 0.0001 |
|  |  |  |  |  |  | Rucker's test statistic/ Cochrane Q-statistics | 0.9961 |
| **Amyotrophic lateral sclerosis (ALS)** | IVW | 4 | 0.995 | 0.784-1.263 | 0.9514 | MR-Egger intecept (p-value) | 0.8013 |
|  | MR-Egger |  | 1,053 | 0.425-2.606 | 0.8297 | I square (IVW) | 0.0000 |
|  | WME |  | 0.998 | 0.915-1.089 | 0.9844 | Cochrane Q-test (IVW) (p-value) | 0.6030 |
|  | MBE |  | 0.952 | 0.775-1.171 | 0.6749 | Rucker's Q-test (p-value) | 0.4118 |
|  |  |  |  |  |  | Rucker's test statistic/ Cochrane Q-statistics | 0.9564 |
| **Age-related macular degeneration (AMD)** | IVW | 38 | 0.989 | 0.945-1.035 | 0.6427 | MR-Egger intecept (p-value) | 0.3854 |
|  | MR-Egger |  | 0.962 | 0.887-1.042 | 0.3300 | I square (IVW) | 0.1454 |
|  | WME |  | 0.976 | 0.947-1.005 | 0.4144 | Cochrane Q-test (IVW) (p-value) | 0.2205 |
|  | MBE |  | 0.981 | 0.918-1.047 | 0.5647 | Rucker's Q-test (p-value) | 0.2139 |
|  |  |  |  |  |  | Rucker's test statistic/ Cochrane Q-statistics | 0.9797 |
| **Multiple sclerosis (MS)** | IVW | 70 | 1,010 | 0.978-1.044 | 0.5289 | MR-Egger intecept (p-value) | 0.8676 |
|  | MR-Egger |  | 1,014 | 0.958-1.074 | 0.6179 | I square (IVW) | 0.0776 |
|  | WME |  | 0.984 | 0.961-1.007 | 0.4817 | Cochrane Q-test (IVW) (p-value) | 0.2954 |
|  | MBE |  | 0.983 | 0.932-1.038 | 0.5438 | Rucker's Q-test (p-value) | 0.2673 |
|  |  |  |  |  |  | Rucker's test statistic/ Cochrane Q-statistics | 0.9998 |

Genetic predisposition to AD was further observed to be a risky for AMD (OR = 1.759, 95% CI = 1.040-1.974, P = 0.0363). On the contrary, AMD was observed to be strongly protective towards MS using MR-Egger method in the presence of significant pleiotropy (MR-Egger intercept p-value = 0.0481, OR = 0.861, 95% CI = 0.776-0.955, P = 0.0059).

## Discussion

The present study is the first study to comprehensively explore causal relationship among various neurodegenerative disorders. My results strongly confirm the genetic relationship between PD, ALS and PD as observed in the case of patients with kii-ALS or Guam-ALS or ALS-PDC. My results further suggest risky and protective effect of AMD towards AD and MS which is consistence with the relative prevalence of retinal degeneration seen in cases of AD and MS (See **Supplementary table 2**).

My study has several strengths and limitations. It is one of the most comprehensive study exploiting the genetics of neurodegenerative disorders to understand the relationship among different disorders. However, differential genomic coverage and different sample sizes of different datasets make it difficult to compare the results. It is quite possible that healthy controls used in different GWAS datasets may be overlapping, leading to the risk of bias in the findings. Nevertheless, I used improved version of IVW method which takes care of these biases. Another limitation could be my inability to explore causality using different types of dementias including Frontotemporal dementia (FTD) which is known to occur in combination with ALS, as it was observed that FTD dataset was highly underpowered with sample size <5000 individuals.

In future, it would be important to dissect different biological pathways using relevant genetic instrument for each pair of relationships. Nevertheless, my study shows utility of genetic data to unearth important biological findings and could enhance our understanding of interconnected etiopathologies of neurodegenerative disorders. Moreover, the finding could impact the diagnosis and management of neurodegenerative disorders.

## Acknowledgement

I thank Prof. Inke König for providing the institutional facilities and research environment for conduct of the research. I acknowledge the investigators of the International Age-related Macular Degeneration Consortium (IAMDGC) for sharing the summary statistics from GWAS on AMD, International Genomics of Alzheimer’s patients (IGAP) on AD, International Parkinson’s Disease Genomics Consortium (IPDGC) for sharing the summary statistics from GWAS on PD, project MinE for sharing the summary statistics from GWAS on ALS, and the International Multiple Sclerosis Genetics Consortium (IMSGC) for sharing the summary statistics on MS.

---

**Supplementary table 1**. Summary statistics used for the conduct of causal analysis in the current manuscript.

**Supplementary table 2.** Examples of recent case reports or case-series reporting co-existence of various neurodegenerative disorders (in alphabetical order).

